# STAT5: A Target of Antagonism by Neurotropic Flaviviruses

**DOI:** 10.1101/606608

**Authors:** Matthew G. Zimmerman, James R. Bowen, Circe E. McDonald, Ellen Young, Ralph S. Baric, Bali Pulendran, Mehul S. Suthar

## Abstract

Flaviviruses are a diverse group of arthropod-borne viruses responsible for numerous significant public health threats; therefore, understanding the interactions between these viruses and the human immune response remains vital. Earlier work has found that WNV and ZIKV infect human DCs and can block antiviral immune responses in DCs. Previously, we used mRNA sequencing and weighted gene co-expression network analysis (WGCNA) to define molecular signatures of antiviral DC responses following activation of innate immune signaling (RIG-I, MDA5, or type I IFN signaling) or infection with WNV. Using this approach, we found that several genes involved in T cell co-signaling and antigen processing were not enriched in DCs during WNV infection. Using cis-regulatory sequence analysis, STAT5 was identified as a regulator of DC activation and immune responses downstream of innate immune signaling that was not activated during either WNV or ZIKV infection. Mechanistically, WNV and ZIKV actively blocked STAT5 phosphorylation downstream of RIG-I, IFNβ, and IL-4, but not GM-CSF signaling. Unexpectedly, dengue virus serotypes 1-4 (DENV1-4) and the yellow fever 17D vaccine strain (YFV-17D) did not antagonize STAT5 phosphorylation. In contrast to WNV, ZIKV inhibited JAK1 and TYK2 phosphorylation following type I IFN treatment, suggesting divergent mechanisms used by these viruses to inhibit STAT5 activation. Combined, these findings identify STAT5 as a target of antagonism by specific pathogenic flaviviruses to subvert the immune response in infected DCs.

**Importance:** Flaviviruses are a diverse group of insect-borne viruses responsible for numerous significant public health threats. Previously, we used a computational biology approach to define molecular signatures of antiviral DC responses following activation of innate immune signaling or infection with WNV. In this work, we identify STAT5 as a regulator of DC activation and antiviral immune responses downstream of innate immune signaling that was not activated during either WNV or ZIKV infection. WNV and ZIKV actively blocked STAT5 phosphorylation downstream of RIG-I, IFNβ, and IL-4, but not GM-CSF signaling. However, other related flaviviruses, dengue virus serotypes 1-4 and yellow fever 17D vaccine strain, did not antagonize STAT5 phosphorylation. Mechanistically, WNV and ZIKV showed differential inhibition of Jak kinases upstream of STAT5, suggesting divergent countermeasures to inhibit STAT5 activation. Combined, these findings identify STAT5 as a target of antagonism by specific pathogenic flaviviruses to subvert antiviral immune responses in human DCs.

## Introduction

West Nile virus (WNV) is a neurotropic flavivirus and the leading cause of arboviral neuroinvasive disease since its introduction to the United States in 1999, accounting for >95% of reported cases (1). After inoculation by an infected mosquito, 20-25% of WNV-infected individuals develop symptoms ranging from mild flu-like symptoms to severe neurologic disease including meningitis, encephalitis, and acute flaccid paralysis (2). Longer-lasting sequelae of WNV-induced fever and neuroinvasive disease include ocular abnormalities, arthralgias, psychological impairment, and permanent memory loss (3). Recently, WNV has also been shown in mice to translocate across the placenta and infect fetal neuronal tissue during pregnancy, prompting fetal demise (4). The lack of FDA-approved antiviral therapeutics or vaccines for human WNV infection reinforces the need to further understand innate immune signaling during WNV pathogenesis.

Dendritic cells (DCs) are vital components of both the innate and adaptive immune responses during flavivirus infection. DCs are professional antigen presenting cells that, upon infection, can process and present antigens on MHC molecules, express co-stimulatory markers, and produce cytokines necessary for activation of the adaptive immune response (5). Mouse models of WNV pathogenesis have demonstrated that DCs are primary targets of infection by WNV and that innate immune signaling in DCs is essential for clearance of neuroinvasive disease (6–9). In the absence of type I interferon (IFN) signaling in murine DCs, WNV shows increased replication in myeloid cells and enhanced tissue tropism, resulting in increased mortality in mice (7). Adoptive transfer studies in mice have also demonstrated that immune competent DCs are essential for proper priming of WNV-specific T cell responses and viral clearance from the central nervous system (8). Recent studies have also established that WNV and a related neurotropic flavivirus, Zika virus (ZIKV), productively infect human monocyte-derived dendritic cells (moDCs) and suppress expression of co-stimulatory markers on infected DCs (10). WNV-infected DCs also display a reduced capacity to induce allogeneic CD4+ and CD8+ T cell proliferation (11). However, the exact mechanism of viral antagonism of human DC activation during WNV infection remains unknown.

During infection in DCs, intracellular WNV RNA is detected by the RIG-I like receptors (RLRs), RIG-I and MDA5, which interact with the adaptor protein MAVS to induce transcription of pro-inflammatory cytokines, antiviral effectors, and type I interferon (IFN) (12, 13). Upon release of type I IFN (IFNα/β), this signal is further potentiated through the type I IFN receptor (IFNAR), causing phosphorylation of bound Janus kinases (JAKs) and subsequent signal transducer and activator of transcription (STAT) proteins (14). Once phosphorylated, STAT proteins will dimerize and translocate to the nucleus, bind to IFN-stimulated response elements (ISREs) on DNA, and initiate robust induction of interferon-stimulated genes (ISGs) to restrict viral replication. Canonically, IFNAR signals through STAT1 and STAT2 heterodimers, both of which are targets of flavivirus antagonism to inhibit antiviral responses (10, 15-18).

In addition, IFNAR can signal through STAT5, a pleiotropic STAT protein activated downstream of numerous cytokines and growth factors. STAT5 exists as two homologs, STAT5a and STAT5b, which share 96% homology at the protein level, which homodimerize upon activation and bind gamma-activated sequences (GAS) elements on DNA (19). STAT5 signaling plays a variety of roles involving DC development and activation. Generation of conventional DCs requires STAT5 signaling by inhibition of plasmacytoid DC development through induction of IRF8 (20). Conditional knockout mouse studies have also established that STAT5 signaling is essential for thymic stromal lymphopoietin (TSLP)-induced activation of DCs and allergic Th2 responses in the lungs (21). Inhibition of STAT5 signaling in monocyte-derived dendritic cells (moDCs) also inhibits lipopolysaccharide-induced DC maturation, costimulatory marker expression, and stimulation of Th1 immune responses (22). Despite the role of STAT5 in DC development and maturation, the importance of STAT5 signaling in DCs during viral infection is currently unknown.

Here, we employed a co-expression network-based analysis combined with promoter scanning analysis, we identified STAT5, a critical transcription factor for regulating antiviral responses and activation within human DCs, downstream of RLR and type I IFN signaling. WNV and ZIKV infection induced minimal STAT5 signaling, corresponding with a failure to up-regulate innate immune mediators and molecules involved in DC activation. The minimal activation of WNV infected DCs reflected viral antagonism of STAT5, and to a lesser extent STAT1 and STAT2, phosphorylation. WNV and ZIKV antagonism of STAT5 was also receptor-specific, inhibiting type I IFN, IL-4, but not GM-CSF, induced STAT5 phosphorylation downstream of JAK kinase signaling. Surprisingly, DENV1-4 and YFV-17D did not block STAT5 signaling, suggesting that targeting STAT5 may be a virus-specific strategy of WNV and ZIKV to subvert T cell immunity.

## Results

### STAT5 is a regulatory node of antiviral DC responses

Previous work from our lab utilized a systems biology approach to assess the global antiviral response during WNV infection within primary human monocyte-derived DCs. Using moDCs from 5 different donors, we performed messenger RNA sequencing following treatment with innate immune agonists targeting the RIG-I, MDA5, and IFNAR signaling pathways as well as infection with WNV at 12 and 24 hpi, signifying log phase viral growth (11). Using weighted gene co-expression network analysis (WGCNA) and Metacore pathway analysis, we determined that the innate immune agonist treatments and WNV infection at 24 hpi induced notable gene expression within the M5 module, a subset of differentially expressed genes (DEGs) highly enriched for pathways involved in type I IFN signaling, PRR signaling, and innate antiviral responses. WGCNA clusters together DEGs into modules based on co-expression, suggesting the presence of common transcriptional regulators driving gene expression within a module. To define the transcriptional regulatory network of the M5 antiviral module, we performed cis-regulatory sequence analysis to computationally predict regulatory nodes using iRegulon, which identifies enrichment of transcription factor binding motifs within the top highly connected genes comprising M5 (23). Consistent with pathway enrichment for antiviral pathways, our analysis identified the ISGF3 transcription complex, IRF1, and NF-κB within the top predicted transcriptional regulators of M5 following RIG-I stimulation. Unexpectedly, we also found notable enrichment for STAT5, a transcriptional regulator with a previously described role in promoting DC activation (21, 22) (Fig. 1A).

**Fig 1.**
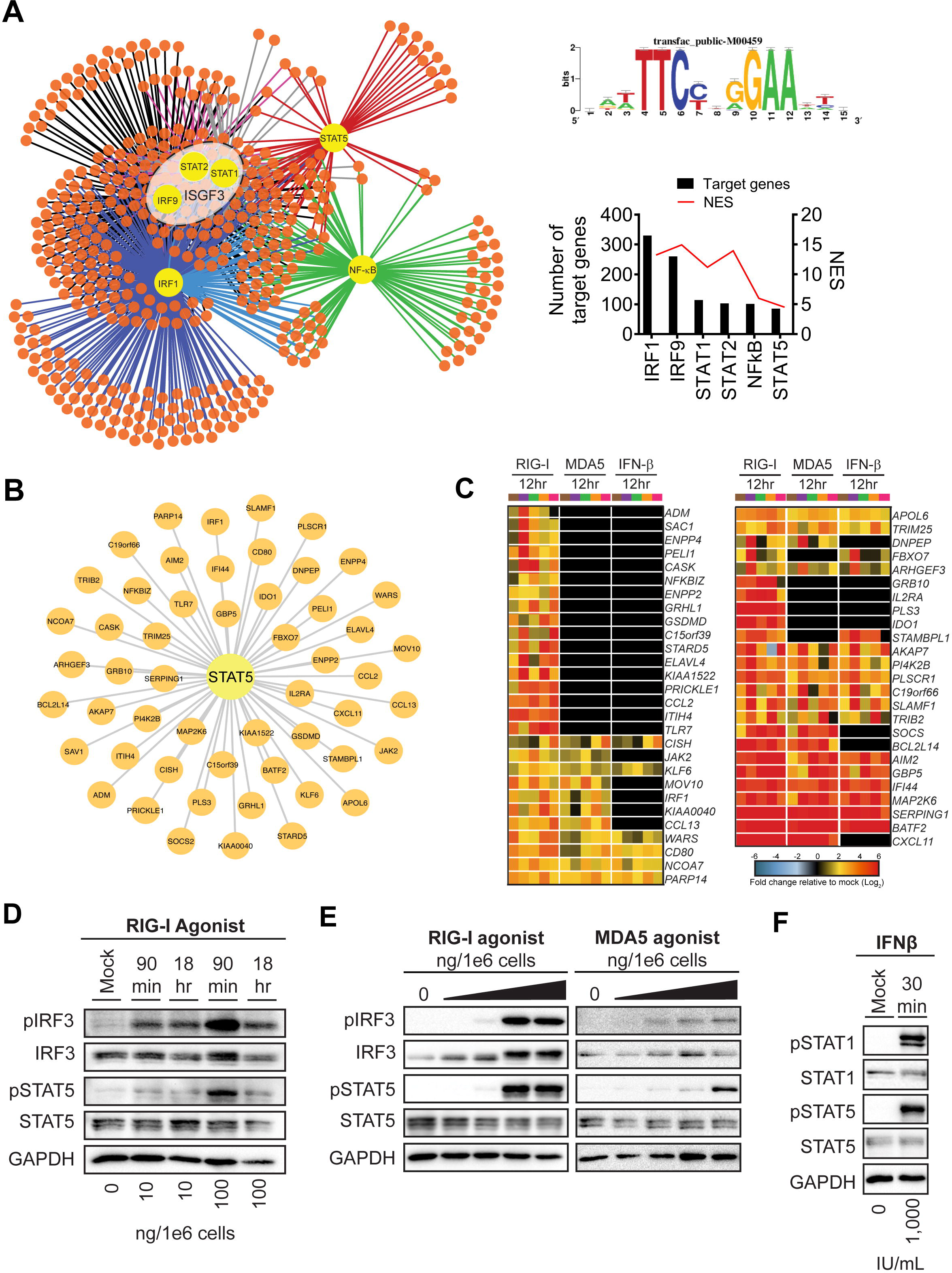
Systems biology reveals STAT5 as a regulatory node of antiviral DC responses. **(A)** Transcription factor regulatory network of module 5 gene expression as predicted by iRegulon (left panel). The top predicted transcriptional regulators (large, yellow nodes) are shown with a connecting line to predicted target genes (small, orange nodes). The consensus sequence for promoter regions targeted by STAT5 as generated by iRegulon and annotated as transfac_public-M00459 (top right panel). The number of predicted target genes and the normalized enrichment score (NES) for a given regulator is shown below (bottom right panel). **(B)** STAT5 regulatory node (large, central node) is shown with the predicted target genes indicated (small nodes), as determined by iRegulon. **(C)** Heatmap of predicted STAT5 target genes after stimulation with innate immune agonists with the log_2_ normalized fold change relative to uninfected, untreated cells is shown (>2-fold change, significance of p<0.01). Genes that did not reach the significance threshold are depicted in black color. Each column within a treatment condition is marked by a unique color and represents a different donor (n= 5 donors). **(D)** moDCs were treated with RIG-I agonist (10 or 100ng/1e6 cells) for 90 minutes or 18hrs. **(E)** moDCs were treated with RIG-I or MDA5 agonist for 90min (10, 100, 1,000, and 10,000ng/1e6 cells). **(F)** moDCs were treated with IFNβ (1000 IU/mL) for 30 minutes. For D-F, Western blot analysis was performed for the indicated proteins. Western blots are representative of data obtained from 3-8 donors.

### STAT5 regulates expression of genes associated with innate immunity and DC activation

Given that a role for STAT5 has not been previously implicated during flavivirus infection, we next evaluated the expression levels of predicted STAT5 target genes. Predicted STAT5 target genes included multiple genes associated with innate immunity (e.g. IRF1, TLR7, TRIM25) and DC activation (e.g. CD80, CXCL11, CCL2) (Fig. 1B). RIG-I agonist induced up-regulation of predicted STAT5 target genes. Transfected poly(I:C) and type I IFN, although to a lesser extent than RIG-I agonist treatment, also activated transcription of several STAT5 target genes. Given recent work implicating STAT5 signaling upstream of DC activation, in part through binding to the promoter regions of CD80 and CD83, we hypothesized that STAT5 might be an important regulator of DC activation downstream of RLR signaling (21, 22). Indeed, multiple predicted STAT5 target genes were involved in processes related to DC activation, including molecules involved in T cell co-signaling (e.g. *CD80*, *IDO1*, *SLAMF1*) and cytokine signaling (e.g. *CCL2*, *CCL13*, *CXCL11*, IL2RA, *JAK2, SOCS, CISH*) (Fig. 1C). Treatment with RIG-I or MDA5 agonist induced significant up-regulation of STAT5 targets involved in DC activation, corresponding with dose-dependent phosphorylation of STAT5 at tyrosine residue 694, a critical event for STAT5 dimerization and DNA binding, following treatment with RIG-I or MDA5 agonist (24). Notably, STAT5 phosphorylation coincided with the kinetics and magnitude of IRF3 phosphorylation (Fig. 1D-E). IFNβ signaling also promoted STAT5 phosphorylation, confirming previous work describing the activation of STAT5 by type I IFN signaling in other cell types (Fig. 1F) (25–27). Collectively, cis-regulatory analysis revealed STAT5 as a transcriptional regulator of numerous antiviral genes within human moDCs and implicates STAT5 as a regulator of DC responses downstream of the RLR and type I IFN signaling axes.

### WNV and ZIKV actively antagonize STAT5 activation

In contrast to RLR and type I IFN stimulation, STAT5 signaling was substantially less enriched during WNV infection, where almost 80% of predicted STAT5 target genes, including those involved in DC activation, were not significantly expressed over mock-infected cells (Fig. 2A). The minimal induction of STAT5 target genes corresponded to a lack of STAT5 phosphorylation following WNV infection during (24hpi) and after (48hpi) log phase viral growth, despite increased amounts of STAT5 total protein by 48hpi (Fig. 2B-C). Given that WNV infection induces type I IFN secretion 48 hpi (Zimmerman et al. 2019) and the selective lack of STAT5 phosphorylation during WNV infection of moDCs, we hypothesized that WNV antagonizes type I IFN-mediated STAT5 signaling. Indeed, WNV infection potently blocked STAT5 phosphorylation following both RIG-I stimulation and IFNβ treatment at 24 and 48 hpi (Fig. 2B-C). Infection with UV-WNV failed to block RIG-I induced STAT5 phosphorylation, suggesting viral replication is required for inhibition of STAT5 signaling. In notable contrast to STAT5, both STAT1 and STAT2 were phosphorylated during WNV infection, suggesting that STAT5 may be differentially modulated by WNV. ZIKV, a closely related neurotropic flavivirus that productively infects human moDCs (10) also antagonized STAT5 phosphorylation downstream of RIG-I and type I IFN signaling (Fig. 2D). Combined, our data strongly suggest that STAT5 is a target of antagonism by both WNV and ZIKV in human DCs.

**Fig 2.**
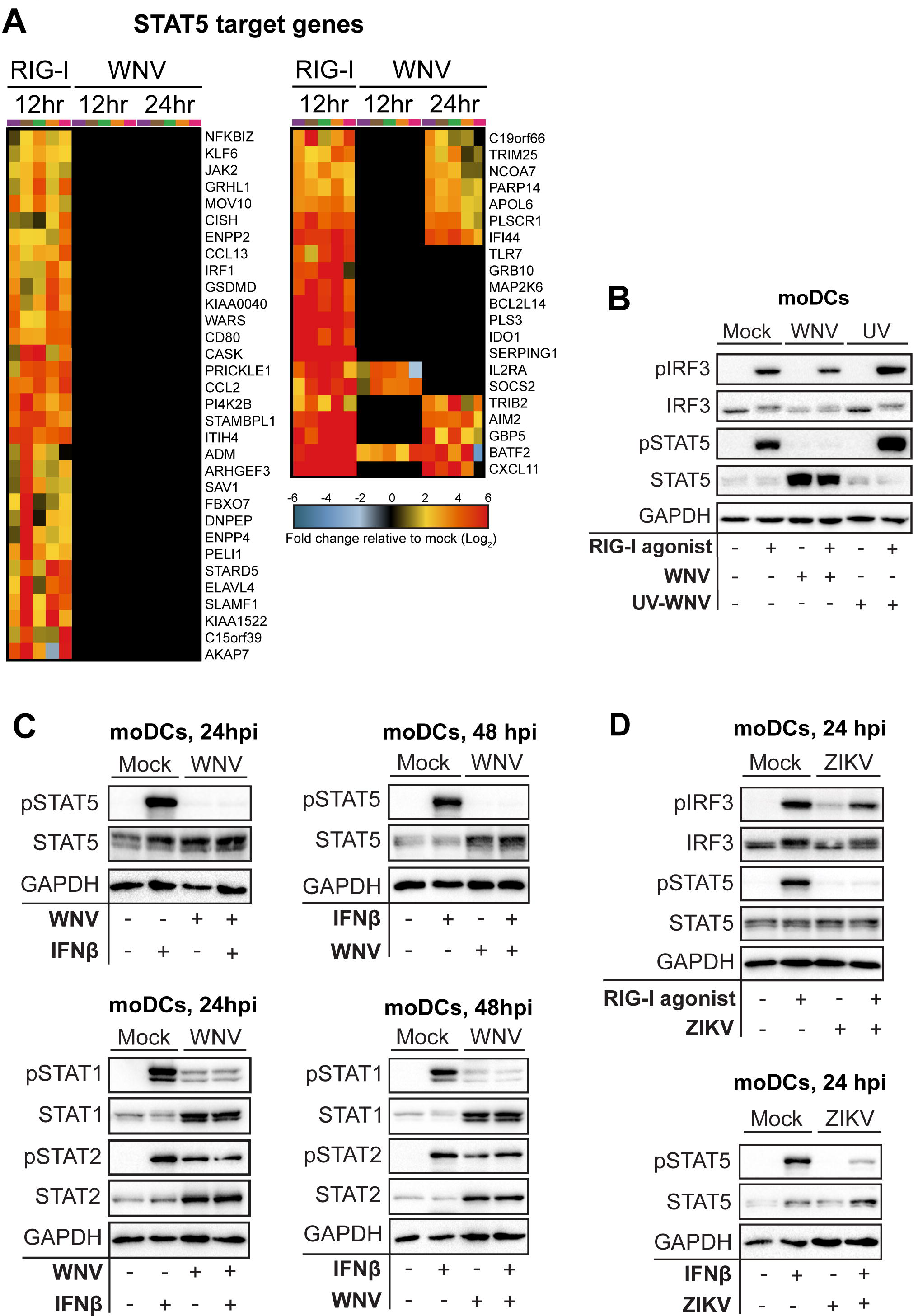
WNV actively blocks STAT5 phosphorylation and expression of STAT5 target genes. **(A)** Heatmap of predicted STAT5 target genes after stimulation with RIG-I agonist or infection with WNV (12, 24 hpi) with the log_2_ normalized fold change relative to uninfected, untreated cells is shown (>2-fold change, significance of p<0.01; right panel). Genes that did not reach the significance threshold are depicted in black color. Each column within a treatment condition is marked by a unique color and represents a different donor (n= 5 donors). **(B)** moDCs were treated with RIG-I agonist treatment (100ng/1e6 cells) for 90 min following no infection (“Mock”), infection with UV-inactivated WNV (MOI 10, “UV-WNV”), or infection with replication competent WNV (MOI 10, “WNV”). **(C, D)** Human moDCs were left uninfected (“Mock”) or infected with WNV or ZIKV (24 or 48hrs, MOI 10 based on Vero titer). Cells were left untreated or pulse-treated with IFNβ (1,000 IU/mL) for 30min. Data in B is representative of results obtained from two independent experiments. Data from C and D representative of results obtained from 3-8 donors.

### WNV and ZIKV block STAT5 phosphorylation in a pathway specific manner

Next, we asked if WNV and ZIKV blocked STAT5 activation downstream of additional cytokine signaling pathways. Common gamma-chain family cytokines, such as IL-4, as well as multiple growth factors, including GM-CSF, signal through their respective receptors to promote STAT5 phosphorylation (28, 29) (Fig. 3A). Similar to our findings with type I IFN signaling, WNV infection dampened IL-4 induced STAT5 phosphorylation in moDCs (Fig. 3B). In contrast, WNV failed to antagonize GM-CSF signaling, where the increased STAT5 protein induced during infection led to increased STAT5 phosphorylation. ZIKV also potently inhibited IL-4, but not GM-CSF, mediated STAT5 phosphorylation in human DCs (Fig. 3C). Together, our findings suggest that WNV blocks STAT5 phosphorylation to antagonize STAT5-dependent gene induction in human moDCs in a pathway specific manner.

**Fig 3.**
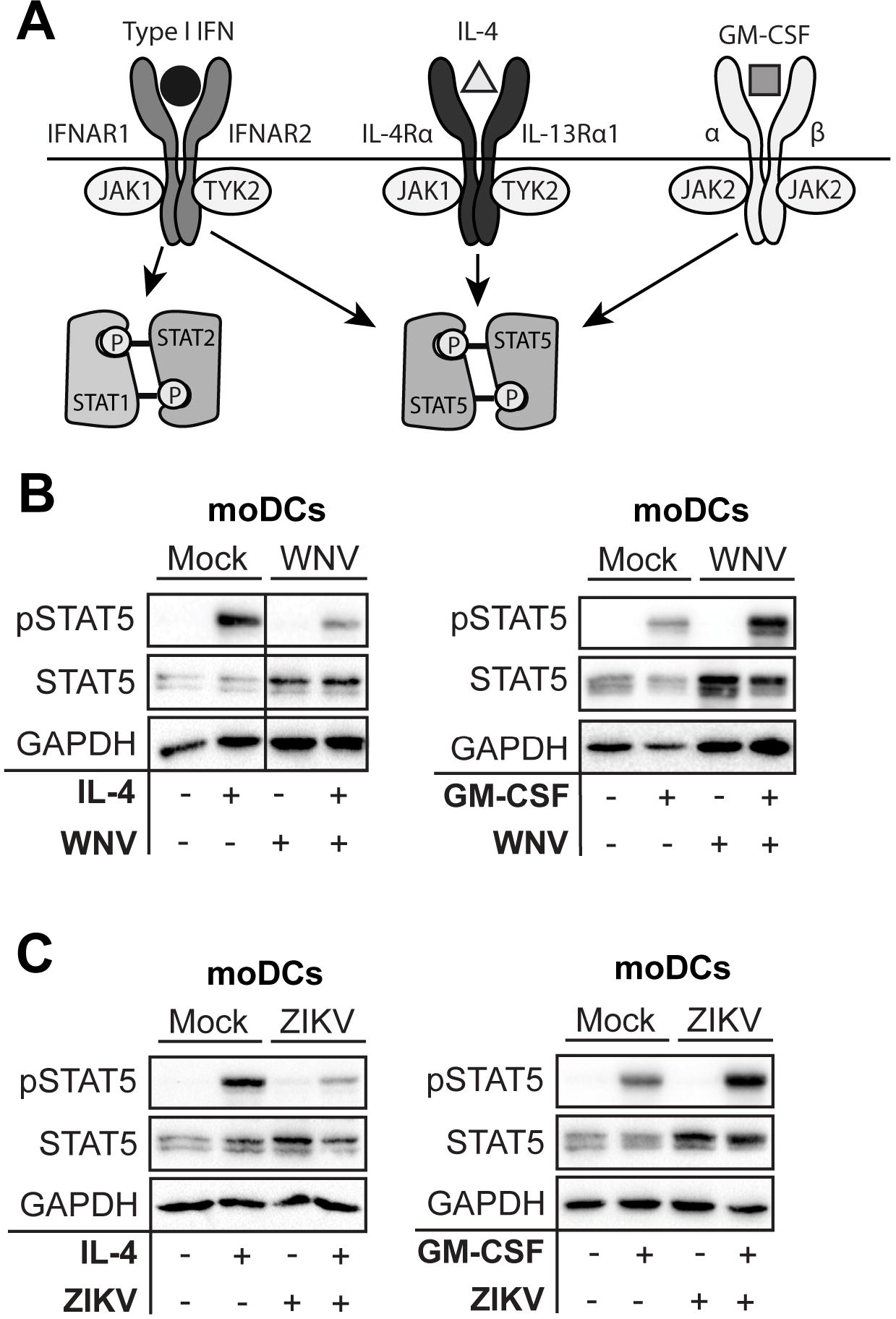
WNV and ZIKV inhibit STAT5 phosphorylation in a pathway-dependent manner. **(A)** Schematic of STAT signaling downstream of type I IFN, IL-4, and GM-CSF signaling. **(B, C)** moDCs were infected with WNV or ZIKV (MOI 10 based on Vero cell titer) for 48hrs and treated with IL-4 (10 ng/mL) for 30 min or GM-CSF (10 ng/mL) for 30min. For B and C, Western blot analysis was performed for the indicated proteins.

### Blockade of STAT5 phosphorylation is flavivirus-specific

Secretion of type I IFN by infected moDCs can also lead to induction of negative regulators that may downregulate type I IFN signaling independent of viral antagonism of STAT proteins. To remove confounding interpretations inherent with IFN-competent cells, we employed a Vero cell model of WNV infection, which allows for synchronous infection of cells that lack endogenous type I IFN signaling. Infection of Vero cells with WNV did not induce phosphorylation of STAT1, STAT2, or STAT5, consistent with the lack of an endogenous type I IFN response. Similar to our studies in human moDCs, pulse treatment of WNV infected Vero cells with IFNβ revealed a substantial blockade of STAT5 phosphorylation (Fig. 4A). STAT1 and STAT2 phosphorylation were also blocked, but in contrast to STAT5, the blockade was of STAT2 was less pronounced. Blockade of STAT5 phosphorylation paralleled the large increase in WNV E and NS3 protein expression between 18 and 24 hpi seen during log phase viral growth. Together, this strongly suggest that antagonism of STAT5 is an active immune evasion mechanism in moDCs and not a bystander effect of endogenous type I IFN signaling during WNV infection. We next asked whether STAT5 antagonism was unique to neurotropic flaviviruses (WNV and ZIKV) or if this mechanism is conserved among other flaviviruses. We infected Vero cells with ZIKV PR-2015, DENV serotypes 1-4, or YFV-17D at an MOI that achieved comparable levels of infection as our studies with WNV (Fig. 4B). Similar to WNV, ZIKV also antagonized STAT5 phosphorylation downstream of type I IFN signaling in Vero cells (Fig. 4C). Consistent with recent work on STAT antagonism by ZIKV, we also observed diminished STAT1 phosphorylation, as well as decreased STAT2 phosphorylation that corresponded with degradation of STAT2 total protein (10, 16). In contrast to WNV and ZIKV, STAT5 phosphorylation was not blocked in Vero cells infected with DENV1-4 and YFV-17D following IFNβ pulse treatment (Fig. 4D-H). Consistent with previous studies of DENV2 antagonism of type I IFN signaling (17), we observed STAT2 degradation in Vero cells infected with DENV1, 2, and 4, while STAT1 phosphorylation was detected after IFNβ pulse treatment. Markedly, DENV3 did not show degradation of STAT2 but showed a slight reduction in STAT2 phosphorylation, suggesting that antagonism of STAT2 by DENV may be a serotype-specific effect. Altogether, these findings demonstrate that antagonism of STAT5 is a virus-specific strategy used by WNV and ZIKV to subvert the antiviral landscape in human DCs during infection.

**Fig 4.**
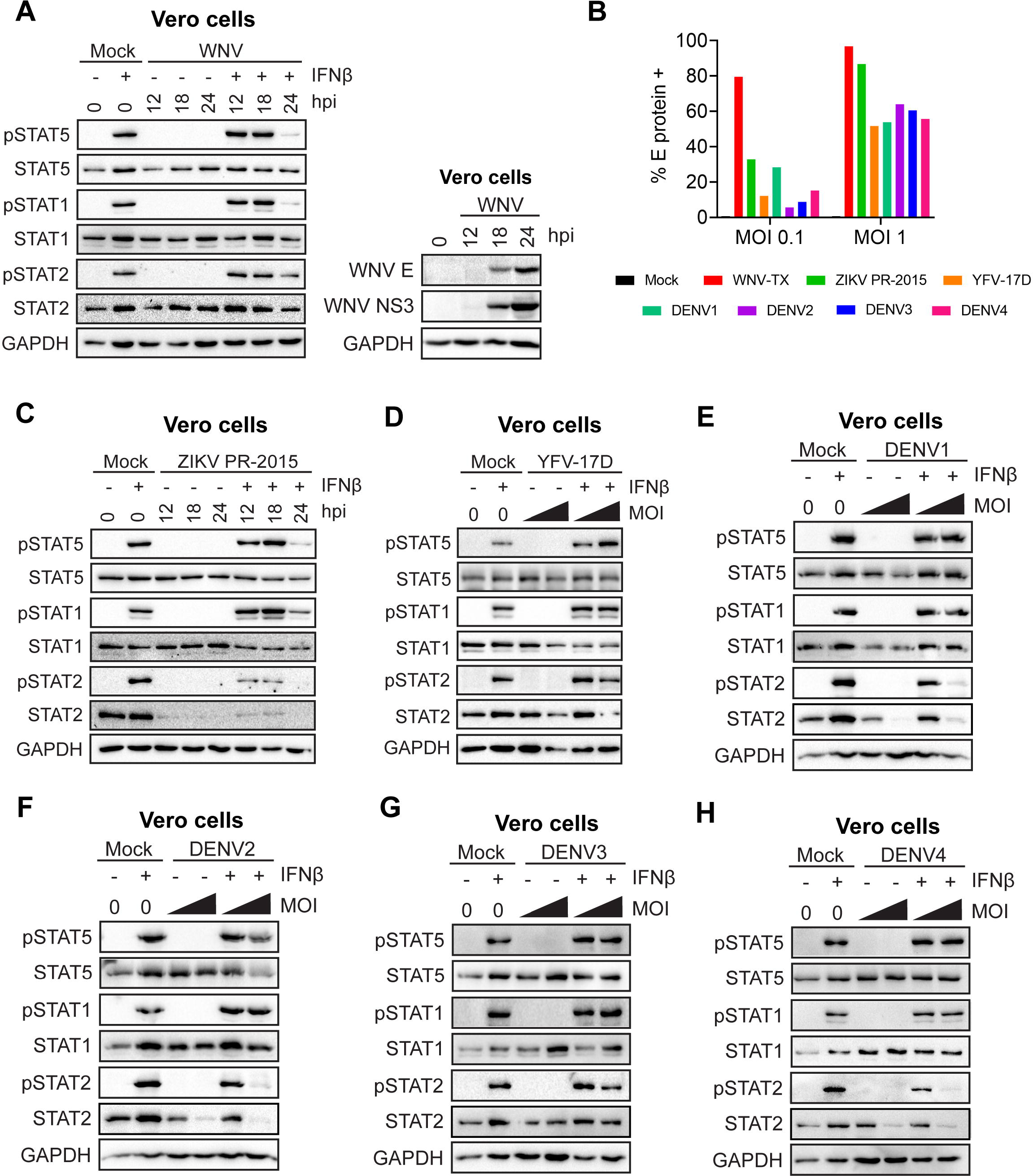
WNV and ZIKV, but not DENV1-4 and YFV-17D, antagonize STAT5 phosphorylation in the absence of IFN signaling. **(A)** Vero cells were left uninfected (“Mock”) or infected with WNV-TX (MOI 1 based on Vero titer) **(B)** Vero cells were infected with WNV-TX, ZIKV-PRVABC59, YFV-17D, or DENV2 (MOI 0.1, 1 based on Vero titer) and percent E protein+ cells were determined by flow cytometry **(C-H)**. Vero cells were left uninfected (“Mock”) or infected with ZIKV (MOI 1 based on Vero titer) **(C)**, YFV-17D (MOI 0.1, 1 based on Vero titer) **(D)**, or DENV1-4 (MOI 0.1, 1 based on Vero titer) **(E-H)** for 24hrs. Cells were then left untreated, or pulse treated with IFNβ (1,000 IU/mL) for 30min. For A and C-H, Western blot analysis was performed for the indicated proteins. Data in C-H is representative of results obtained from two independent experiments.

**Fig 5.**
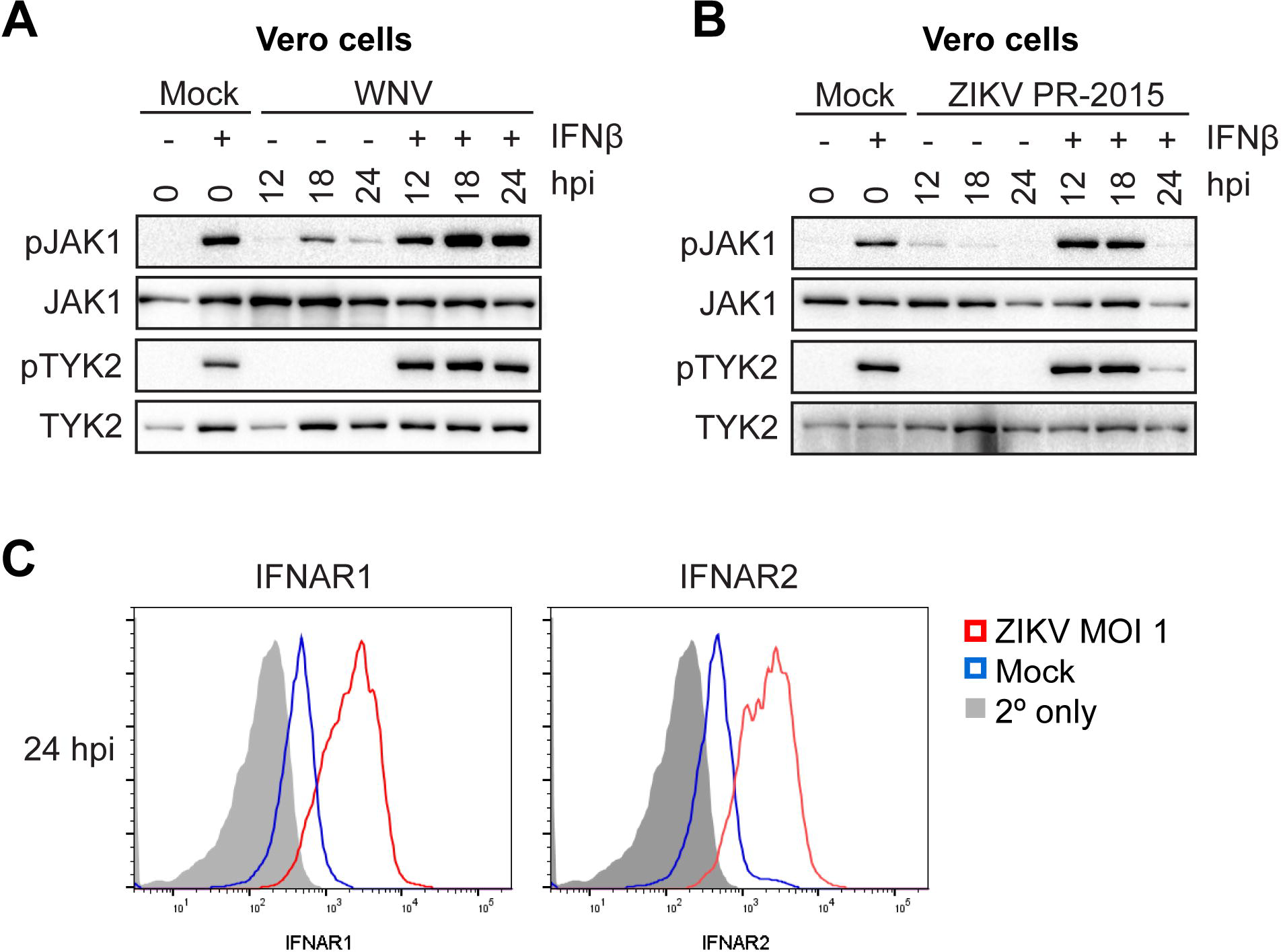
ZIKV, but not WNV, blocks Tyk2 or JAK1 activation to compromise STAT5 phosphorylation. **(A, B)** Vero cells were left uninfected (“Mock”) or infected with WNV (MOI 0.1 based on Vero titer) or ZIKV-PRVABC59 (MOI 1 based on Vero titer) for 24hrs. Cells were then left untreated, or pulse treated with IFNβ (1,000 IU/mL) for 30min. **(C)** Vero cells were left uninfected (“Mock”) or infected with WNV (MOI 1 based on Vero titer) for 24hrs and levels of surface IFNAR1 and IFNAR2 expression were measured by flow cytometry. For A-B, Western blot analysis was performed for the indicated proteins. Data in A and B is representative of results obtained from two independent experiments.

### ZIKV, but not WNV, blocks activation of Tyk2 and JAK1

The pathway specific inhibition of STAT5 through type I IFN, IL-4, but not GM-CSF, signaling within moDCs provides insight into the host target of viral antagonism. The type I IFN receptor and the type II IL-4 receptor, associate with JAK1 and TYK2 to mediate tyrosine phosphorylation of STAT1 and STAT2, while TYK2 constitutively associates with STAT5 and mediates its tyrosine phosphorylation (27, 28). GM-CSF signaling through the GM-CSF receptor predominately activates JAK2, but not JAK1 or TYK2, suggesting TYK2 and JAK1 may be targeted to block STAT5 phosphorylation (29, 30). To assess JAK inhibition, we infected Vero cells at MOI 0.1 with WNV for 12, 18, and 24 hours. Consistent with the lack of endogenous type I IFN production, WNV infection alone did not induce TYK2 phosphorylation. Although we did observe JAK1 phosphorylation, this is likely explained by the production of other cytokines that can activate JAK1. In contrast to blockade of STAT protein phosphorylation, we observed no blockade of TYK2 and enhanced JAK1 phosphorylation following IFNβ pulse treatment of infected cells **(Fig. 7A)**. Despite its similarities to WNV in inhibiting STAT5 phosphorylation, ZIKV efficiently blocked phosphorylation of both JAK1 and Tyk2 downstream of type I IFN signaling **(Fig. 7B)**. Previous work has shown that flaviviruses are capable of downregulating IFNAR1 from the cell surface (31, 32), so, we assessed whether ZIKV infection decreased expression of IFNAR1 or IFNAR2 from the Vero cell surface **(Fig. 7C)**. At 24 hpi where we observed blockade of JAK1 and Tyk2 phosphorylation in ZIKV-infected Vero cells, we observed an increase in both IFNAR1 and IFNAR2 expression compared to mock infected cells. Combined, these findings suggest that WNV and ZIKV antagonize STAT5 signaling through two separate mechanisms: ZIKV at the level of JAK kinase phosphorylation and WNV directly inhibiting phosphorylation of STAT5.

## Discussion

In this study, we combined traditional virologic and immunologic measures with transcriptomic and computational approaches to define the global antiviral response during WNV infection in human primary cells. Using cis-regulatory sequence analysis, STAT5, a transcription factor previously described as a regulator of DC activation, was identified as an important regulatory node of antiviral DC responses downstream of innate immune signaling. In contrast, STAT5 signaling was minimally activated during WNV infection in human moDCs, corresponding with minimal expression of genes of inflammatory responses or molecules involved in T cell priming. Mechanistically, WNV and ZIKV blocked STAT5 phosphorylation downstream of RIG-I, IFNβ, IL-4, but not GM-CSF signaling, suggesting pathway-specific antagonism of STAT5 activation. Notably, neither the related flaviviruses DENV1-4 nor vaccine strain YFV-17D inhibited STAT5 phosphorylation. Mechanistically, WNV and ZIKV displayed differential inhibition of JAK1 and TYK2 phosphorylation, indicating two distinct mechanisms to antagonize STAT5 signaling. Combined, our study identifies antagonism of STAT5 phosphorylation as a conserved immune countermeasure used by certain pathogenic flaviviruses.

Cis-regulatory sequence analysis revealed STAT5 as a regulatory node of multiple components of DC activation downstream of RLR and type I IFN signaling. Indeed, we observed significant enrichment for multiple STAT5 target genes involved in DC activation, which corresponded with increased gene expression and secretion of pro-inflammatory cytokines and up-regulation of proteins involved in T cell activation (11). While this confirms previous studies that have implicated STAT5 upstream of DC activation, our work reveals a new facet of STAT5 activation during flavivirus infection through engagement of innate immune signaling (21, 22). The rapid kinetics of STAT5 phosphorylation following RIG-I or MDA5 stimulation suggest that RLR signaling may directly induce phosphorylation of STAT5 through activation of a tyrosine kinase, such as Src or Lyn, both of which are induced by RLR signaling (33–37). Alternatively, rapid production of type I IFN, which also promotes STAT5 activation, may mediate STAT5 phosphorylation following RLR signaling. Combined, our data suggests that STAT5 is an important regulatory node downstream of the RLR and type I IFN signaling axis.

In contrast to RLR and IFNβ signaling, WNV infection did not up-regulate most predicted STAT5 target genes, and STAT5 was not phosphorylated during infection despite secretion of IFNβ and IFNα. Corresponding with minimal STAT5 enrichment, WNV infection failed to promote up-regulation of inflammatory mediators and molecules involved in antigen presentation and T cell co-signaling. These findings are similar to previous work, where WNV infection also failed to induce inflammatory cytokine secretion (38). Infection of moDCs with a non-pathogenic WNV isolate, WNV Kunjin, also induced minimal production of IL-12, despite notable up-regulation of both CD86 and CD40 (39). This suggests that an inability to induce inflammatory cytokine responses may be shared among WNV strains, while pathogenic strains have evolved unique mechanisms to subvert antigen presentation and T cell activation. The lack of activation of WNV-infected human moDCs is also similar to our recent work with ZIKV (10). In contrast to WNV and ZIKV, infection of moDCs with the YFV vaccine strain (YFV-17D) up-regulates multiple inflammatory mediators and surface expression of CD80 and CD86 (40). The ability of YFV-17D to induce strong DC activation may reflect the loss of a viral antagonist during the attenuation process, similar to the ability of WNV Kunjin to induce up-regulation of CD86 and CD40 (39). Alternatively, the ability of YFV-17D to induce DC activation may be an inherent property of certain flaviviruses. Indeed, DENV has also been found to activate inflammatory responses and up-regulate co-stimulatory molecules following infection (41, 42). We demonstrate here that, unlike WNV and ZIKV, YFV-17D and DENV1-4 do not inhibit STAT5 phosphorylation. This raises the possibility that the inability to block STAT5 may explain the activation induced during YFV-17D and DENV infection. Altogether, we demonstrate that WNV, similar to ZIKV, induces minimal DC activation during productive infection, contrasting with both DENV and YFV-17D.

Blockade of STAT5 signaling by WNV was found to be multi-frontal, as multiple STAT5 signaling cytokines induced downstream of RIG-I signaling (GM-CSF, IL-4, IL-15) are not produced during infection with WNV (11)

The lack of GM-CSF secretion may also overcome the need for WNV to block GM-CSF-induced STAT5 phosphorylation. Similar to WNV, ZIKV has also been found to induce minimal secretion of cytokines, including IL-4, GM-CSF, IL-12, IL-15, and RANTES, during infection of human moDCs (10). In contrast, DENV and YFV-17D infection in human moDCs has been reported to induce proinflammatory cytokine expression and secretion, including IL-12 and RANTES (43–46). This robust cytokine response could potentially reflect the inability of DENV and YFV-17D to inhibit STAT5 signaling cytokines. Combined, this suggests that the ability to antagonize STAT5 signaling may be an important feature of WNV and ZIKV pathogenesis.

Surprisingly, viral blockade of STAT5 phosphorylation during WNV, but not ZIKV infection, did not correspond with impaired activation of TYK2 and JAK1, members of the Janus associated kinase (JAK) family that phosphorylate STAT5 downstream of type I IFN, IL-4, but not GM-CSF signaling. Numerous flaviviruses, including the closely related Japanese encephalitis virus (JEV) and Langat virus, have been shown to inhibit TYK2 activation through the NS5 protein to subvert JAK/STAT signaling (18, 47). Nevertheless, our work suggests that inhibition of TYK2 or JAK1 activation is likely not the mechanism used by WNV to block STAT5 phosphorylation. Indeed, STAT1 and STAT2, which are also phosphorylated by TYK2 and JAK1, were not blocked as strongly as STAT5, suggesting viral inhibition may be occurring at a step unique to STAT5 signaling. STAT5 itself is likely not targeted directly, given that GM-CSF induced STAT5 signaling remains intact during WNV infection. One possibility is that WNV may specifically disrupt the interaction between STAT5 and TYK2 or JAK1. WNV infection may also directly induce negative regulators of STAT5 signaling, such as facilitating interactions between STAT5 and the protein tyrosine phosphatase, SHP-2, a well-known regulator of cytosolic STAT5 signaling (48). Another possibility includes induction of the suppressor of cytokine signaling family, which broadly regulate JAK/STAT signaling and can be modulated during flavivirus infection (49). Importantly, UV-inactivated virus, which undergoes cellular binding and entry but is replication incompetent, was unable to inhibit STAT5. This suggests that viral replication is required for STAT5 antagonism, potentially through a secreted viral protein that would affect both infected and uninfected cells. The observation that DENV1-4 and YFV-17D do not block STAT5 phosphorylation provides a valuable tool to further define the viral and host factors within subsets of flaviviruses that mediate STAT5 blockade.

In summary, our systems biology approach identified STAT5 as a regulator of DC activation that is blocked by WNV as a mechanism to subvert DC activation and T cell priming. ZIKV, but not YFV-17D or DENV1-4, also blocked STAT5 signaling, suggesting that viral antagonism of STAT5 may be a common strategy of particular subsets of pathogenic flaviviruses to evade the pressures of host immunity. Our study advances our understanding of how pathogenic flaviviruses subvert antiviral immunity during human infection.

## Materials and Methods

### Ethics statement

Human peripheral blood mononuclear cells (PBMCs) were obtained from de-identified healthy adult blood donors and processed immediately. All individuals who participated in this study provided informed consent in writing in accordance to the protocol approved by the Institutional Review Board of Emory University, IRB#00045821, entitled “Phlebotomy of healthy adults for the purpose of evaluation and validation of immune response assays”.

### Cell lines

Vero cells (WHO Reference Cell Banks) were maintained in complete DMEM. Complete DMEM was prepared as follows: DMEM medium (Corning) supplemented with 10% fetal bovine serum (Optima, Atlanta Biologics), 2mM L-Glutamine (Corning), 1mM HEPES (Corning), 1mM sodium pyruvate (Corning), 1× MEM Non-essential Amino Acids (Corning), and 1× Antibiotics/Antimycotics (Corning). Complete RPMI was prepared as follows: cRPMI; RPMI 1640 medium (Corning) supplemented with 10% fetal bovine serum (Optima, Atlanta Biologics), 2mM L-Glutamine (Corning), 1mM Sodium Pyruvate (Corning), 1× MEM Non-essential Amino Acids (Corning), and 1× Antibiotics/Antimycotics (Corning).

### Generation of monocyte derived dendritic cells

To generate human moDCs, CD14+ monocytes were differentiated in cRPMI supplemented with 100ng/mL of GM-CSF and IL-4 for 5-6 days, as previously described (10). In brief, freshly isolated PBMCs obtained from healthy donor peripheral blood (lymphocyte separation media; StemCell Technologies) were subjected to CD14+ magnetic bead positive selection using the MojoSort Human CD14 Selection Kit (BioLegend). Purified CD14+ monocytes were cultured in complete RPMI supplemented with 100ng/mL each of recombinant human IL-4 and GM-CSF (PeproTech) at a cell density of 2e6 cells/mL. After 24hr of culture, media and non-adherent cells were removed and replaced with fresh media and cytokines. Suspension cells (“moDCs”) were harvested after 5-6 days of culture and were consistently CD14-, CD11c+, HLA-DR+, DC-SIGN+, and CD1a+ by flow cytometry. For experimentation, moDCs were maintained in complete RPMI without GM-CSF or IL-4. For experiments measuring STAT5 phosphorylation, moDCs were rested in cRPMI without GM-CSF or IL-4 for 24hrs prior to experimentation.

### Viruses

WNV stocks were generated from an infectious clone, WNV isolate TX 2002-HC, and passaged once in Vero cells, as previously described (50). ZIKV strain PRVABC59 was obtained from the Centers for Disease Control and Prevention as previously described (51). YFV −17D was subpassaged from YF-VAX (Aventis Pasteur) in SW-480 cells followed by passaging in Vero cells as previously described (40). The DENV1, DENV3, and DENV4 strains were derived from low passage clinical isolates and generated from infectious clones (ic) as previously described (52–54). The DENV2 ic was generated in the Baric laboratory in a similar manner to the DENV1, DENV3 and DENV4 ic. The generation and characterization of the DENV2 S16803 ic will be described in a future publication. WNV and ZIKV were titrated by plaque assay on Vero cells with a 1% agarose overlay and crystal violet counterstain, as previously described (50). DENV was titrated by focus forming assay as previously described (54). moDCs were infected with WNV or ZIKV at MOI 10 for 1 hr at 37°C in cRPMI (without GM-CSF or IL-4). After 1 hr, virus was washed off and cells were resuspended in fresh cRPMI and incubated at 37°C for 3-72 hr.

### Quantitation of infectious virus

Infectious virus was quantitated using a plaque assay on Vero cells with a 1% agarose overlay and crystal violet counterstain, as previously described (50).

### Innate immune agonists

To stimulate RIG-I signaling, 100ng of RIG-I agonist derived from the 3’-UTR of hepatitis C virus (55) was transfected per 1e6 cells using TransIT-mRNA transfection kit (Mirus). For stimulation of MDA5 signaling, 100ng of high molecular weight poly-(I:C) was transfected per 1e6 cells using LyoVec transfection reagent (Invivogen). To stimulate type I IFN signaling, cells were incubated with 100 IU/mL of human recombinant IFNβ. In select experiments, different doses of agonists were used and this is indicated within the respective Figure legend.

### RNA sequencing and bioinformatics

moDCs were generated from 5 donors and either treated with innate immune agonists for 12hr (RIG-I, MDA5, or IFNβ) or infected with WNV (12hpi and 24hpi). Total RNA was purified (Quick-RNA MiniPrep Kit; Zymo Research) and mRNA sequencing libraries were prepared for RNA sequencing (Illumina TruSeq chemistry). RNA sequencing was performed on an Illumina HiSeq 2500 System (100bp single end reads). Sequencing reads were mapped to the human reference genome 38. Weighted gene co-expression module analysis was performed on DESeq2 normalized mapped reads (TIBCO Spotfire with Integromics Version 7.0) from RIG-I agonist, MDA5 agonist, IFNβ, and mock treated samples. First, the datasets were reduced to focus the network analysis on the 5446 most variable genes (as determined by variation value greater than 1) using the Variance function in R. We constructed a signed weighted correlation network by generating a matrix pairwise correlation between all annotated gene pairs. The resulting biweight mid-correlation matrix was transformed into an adjacency matrix using the soft thresholding power (β1) of 12. The adjacency matrix was used to define the topological overlap matrix (TOM) based on a dissimilarity measurement of 1- TO. Genes were hierarchically clustered using average linkage and modules were assigned using the dynamic tree-cutting algorithm (module eigengenes were merged if the pairwise calculation was larger than 0.75). This resulted in the construction of six modules. Transcriptional regulators within the M5 module were computationally predicted with iRegulon (23), using the top most connected M5 genes using an eigengene-based connectivity cutoff of 0.4. Differentially expressed genes within the M5 module were identified as having a >2-fold change (significance of p<0.01) relative to uninfected and untreated cells. Pathway analysis was performed on M5 genes using MetaCore pathway map analysis (version 6.29, Thomson Reuters). The raw data of all RNA sequencing will be deposited into the Gene Expression Omnibus (GEO) repository and the accession number will be available following acceptance of this manuscript.

### Flow cytometry

Cells were prepared for analysis as previously described (10). In brief, cells were Fc receptor blocked for 10 min, stained for phenotypic and activation markers for 20 min, and viability stained for 20 min (Ghost Dye Violet 510, Tonbo Biosciences). For intracellular staining of WNV E protein, cells were fixed and permeabilized (Transcription Factor Staining Buffer Kit, Tonbo Biosciences) and labeled with E16-APC for 20min at room temperature (56). Flow cytometry data was analyzed using FlowJo version 10 software. ImageStream data was analyzed using the Amnis IDEAS software. Primary antibodies are listed in S1 Table.

### Western blot

Whole-cell lysates were collected in modified radioimmunoprecipitation assay buffer (10 mM Tris [pH 7.5], 150 mM NaCl, 1% sodium deoxycholate, and 1% Triton X-100 supplemented with Halt Protease Inhibitor Cocktail [ThermoFisher] and Halt Phosphatase Inhibitor Cocktail [ThermoFisher]). Protein lysates were separated by SDS-PAGE and western blot analysis was performed using the ChemiDoc XRS+ imaging system (BioRad). Western blots were analyzed using Image Lab version 5.2.1 software (BioRad) and prepared for publication using Adobe Illustrator. Primary antibodies are listed in S1 Table.

### Statistics

All statistical analysis was performed using GraphPad Prism version 8 software. The number of donors varied by experiment and is indicated within the Figure legends. Statistical significance was determined as P<0.05 using a Kruskal-Wallis test (when comparing more than two groups lacking paired measurements), a Wilcoxon test (when comparing two groups with paired measurements. All comparisons were made between treatment or infection conditions with a time point matched, uninfected and untreated control.

## Acknowledgements

We thank Aaron Brault for his help and advice on working with the Zika virus isolate PRVABC59, the Children’s Healthcare of Atlanta and Emory University Pediatric Flow Cytometry Core for providing access to flow cytometry, ImageStream, and luminex systems, and the Yerkes Genomics Core for performing RNA sequencing.

